# Redescription of *Dendrophthoe falcata* (L. f.) Ettingsh (Loranthaceae) with notes on haustorium

**DOI:** 10.1101/2023.09.29.560265

**Authors:** Somnath Bhakat

## Abstract

Morphology of *Dendrophthoe falcata* (L. f.) Ettingsh var coccinea is described in detail from West Bengal with notes on the histology of haustorium. Structural peculiarity of the style is explained in the light of embryo sac development. Section of immature fruit shows different zones including embryo.

Three types of haustoria namely woody gall, clasping union ad epicortical root is observed in different species of the host plant. Section of haustorium shows penetration peg which reach the host xylem to suck the sap. In guava plant, two types of haustoria develop – primary or true haustorium which penetrate the host stem and secondary haustorium which develop in between two roots of the parasite. Secondary haustorium is oval in shape with a few concentric rings like thickening and never develops penetration peg.

## Introduction

Honey suckles mistletoe, *Dendrophthoe falcata* (L.f.) Ettingsh is an aerial hemiparasitic epiphyte of woody plant. It is known as “Vanda” in the Indian Ayurvedic system of Medicine. The mistletoe is a chlorophyllus photosynthetic stem parasite that connect to vascular tissues of host stem for water and nutrients (Mathiasen et al. 2008). They cause severe damage to many economically important plants like mango (*Mangifera indica*), guava (*Psidium guajava*), sugar apple (*Annona squamosa*) etc. The genus *Dendrophthoe* Mart. comprises about 30 species distributed in tropical Africa, Asia, and Australia (Nickrent et al. 2010, Mabberley 2017) of which eight species are reported in India (Singh et al. 2020b, Sivaramkrishna et al. 2021). *D. falcata* is one of the eight species that is very common in Indian sub-continent. The plant is reported from almost all states of India including West Bengal. So far, it has been reported to exist in more than 300 host plants (Sampathkumar and Selvaraj 1981). Two of its varieties are distributed in India, namely, var. falcata (honey suckle mistletoe) with white flowers and var. coccinea (red honey suckle mistletoe) with red flowers.

Among different species of *Dendrophthoe, D. falcata* is largely studied, mostly for its medicinal importance (Sastry 1952, Pattanayak and Sunita 2008, Pattanayak et al. 2008) and a few on other aspects like morphology, host plant etc. (Singh, 1952; Sampathkumar and Selvaraj, 1981). But reports on description of the whole plant including its floral structure and haustorium is scanty. The present paper describes the morphology of whole plant including the floral structure in detail with notes on the structure of haustorium of *Dendrophthoe falcata* var. coccinea.

## Materials and methods

Field explorations were conducted in district Birbhum (23°32’30’’N to 24°35’00’’N, 87°05’25’’E to 88°01’40’’E), West Bengal, India. Herbarium specimens were prepared using standard technique. Leaves and flowers were preserved in FAA. The morphological descriptions and illustrations were based on fresh specimens and field collection. I observed the red mistletoe in huge number of mango plants (*Mangifera indica*), in a very few guavas plant (*Psidium guajava*), a citrus plant (*Citrus maxima*), a wood apple plant (*Aegle marmelos*), a fig tree (*Ficus carica*) and a flowering plant (*Cascabela thebetia*). Parasites in the host plant were photographed in the field. To study the anatomy of the haustorium of *D. falcata*, a cross sectional cut with a handsaw at the attachment point of the mistletoe was done. Then the dried wood samples were sanded with fine sandpaper and after removing the dust, photos of different parts were taken carefully.

From a mature bud, through a small incision on the upper part of the throat, pollen grains were collected from anthers in a small watch glass, washed by alcohol, then mounted in a thin glass slide by 50% glycerine. Pollen grains were viewed under stereo microscope and measured by ocular micrometer.

Section of ovary and style (of the lower part of the bend) was prepared and observed under microscope. Section of an immature fruit was also made to find out tissue structure of the fruit, especially the viscid layer. The mature fruit is squeezed from the anterior end, the seed along with a white viscid mucilage came out of the fruit from posterior end. After washing the mucilage in alcohol, the seed was measured. To get the embryo intact, mesocarp and endosperm is removed carefully one after another with the help of needle and blade.

I have collected growing root tip of *D. falcata* from guava plant (Fig. x) and sectioned it to observe the distribution of vascular bundle. In the same plant a root of the parasite folded and the two parts run parallel to each other with three haustoria-like attachment (Fig. x). I have separated the two parts and photographed to find out the nature of attachment.

Identity of the specimens were confirmed by scrutinizing the relevant literature (Danser, 1929; Johri and Bhatnagar,1972; Rajsekharan, 2012; Singh et al., 2020a; Singh, 1952). A comparative account of eight Indian species of the genus *Dendrophthoe* was prepared using data from different sources.

## Result

An evergreen, large, woody, hemiparasitic shrub mostly on the branches of large Mango trees (*Mangifera indica* L. family Anacardiaceae) (Fig. 3).

**Fig, 1.**
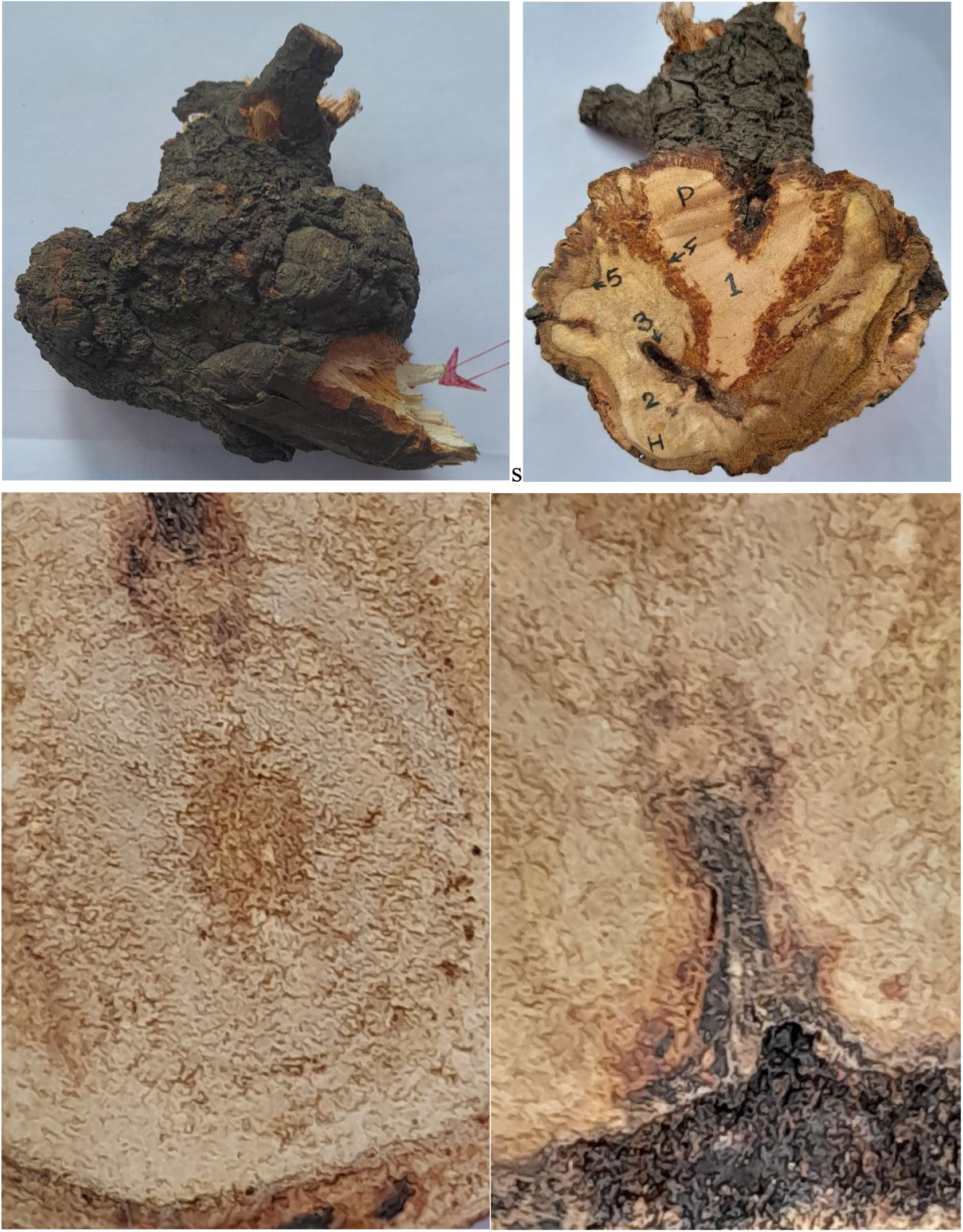
*Dendrophthoe falcata* (Clockwise from top left): A haustorium (arrow indicate host stem); section of haustorium H-host, P-parasite, 1-penetration peg, 2-host cambial zone, 3-collapsed zone, 4-reddish brown secretion, 5-bark like dermal tissue of haustorium; xylem connective in between host and parasite; host tissue with central core.

**Fig. 2.**
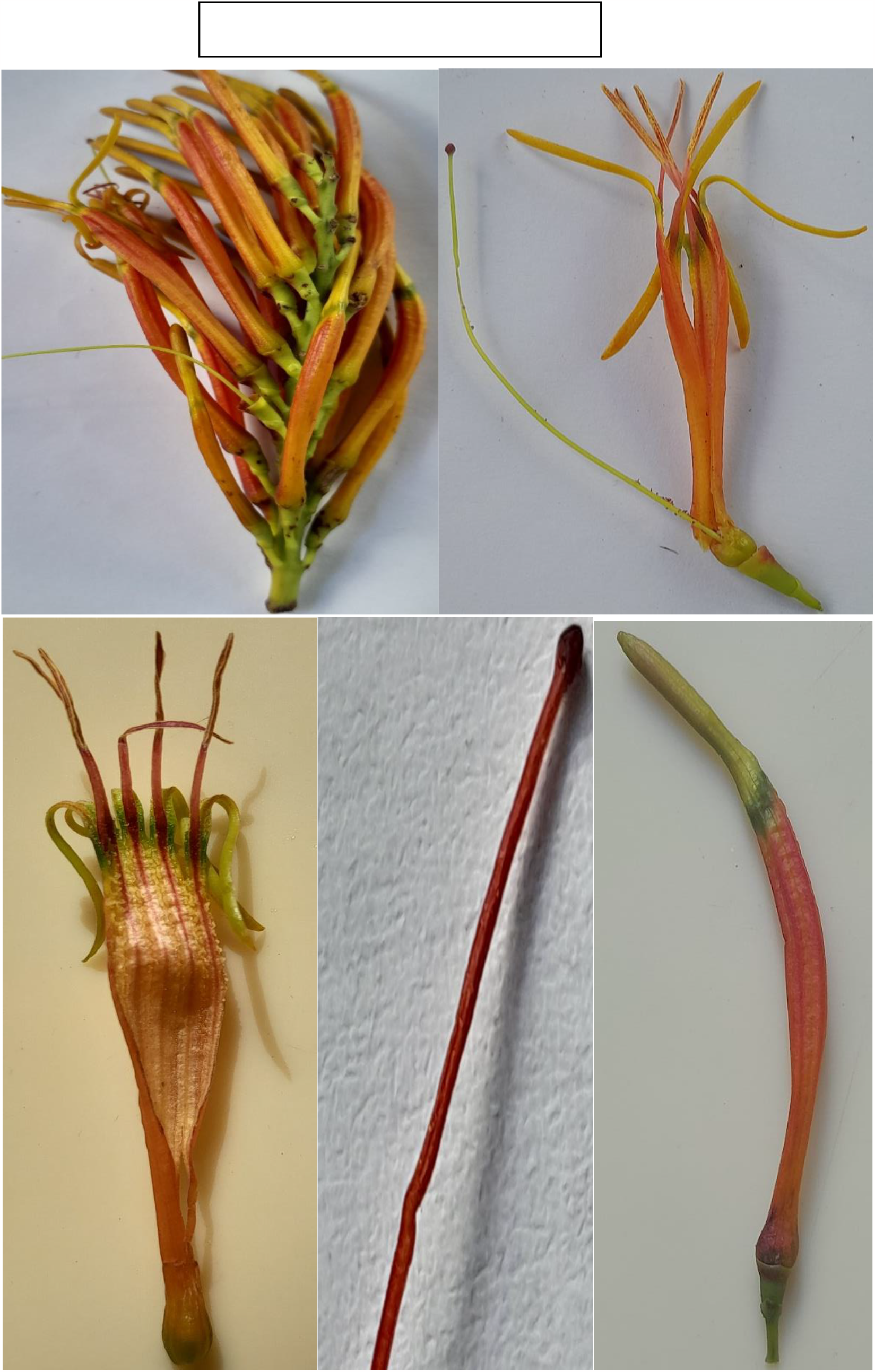
*Dendrophthoe falcata* (Clockwise from top left): A bunch of bud; dissected flower showing stamen and style; a mature flower bud; style with distinct bend; attachment of filament with corolla, and free lobes of corolla.

**Fig. 3.**
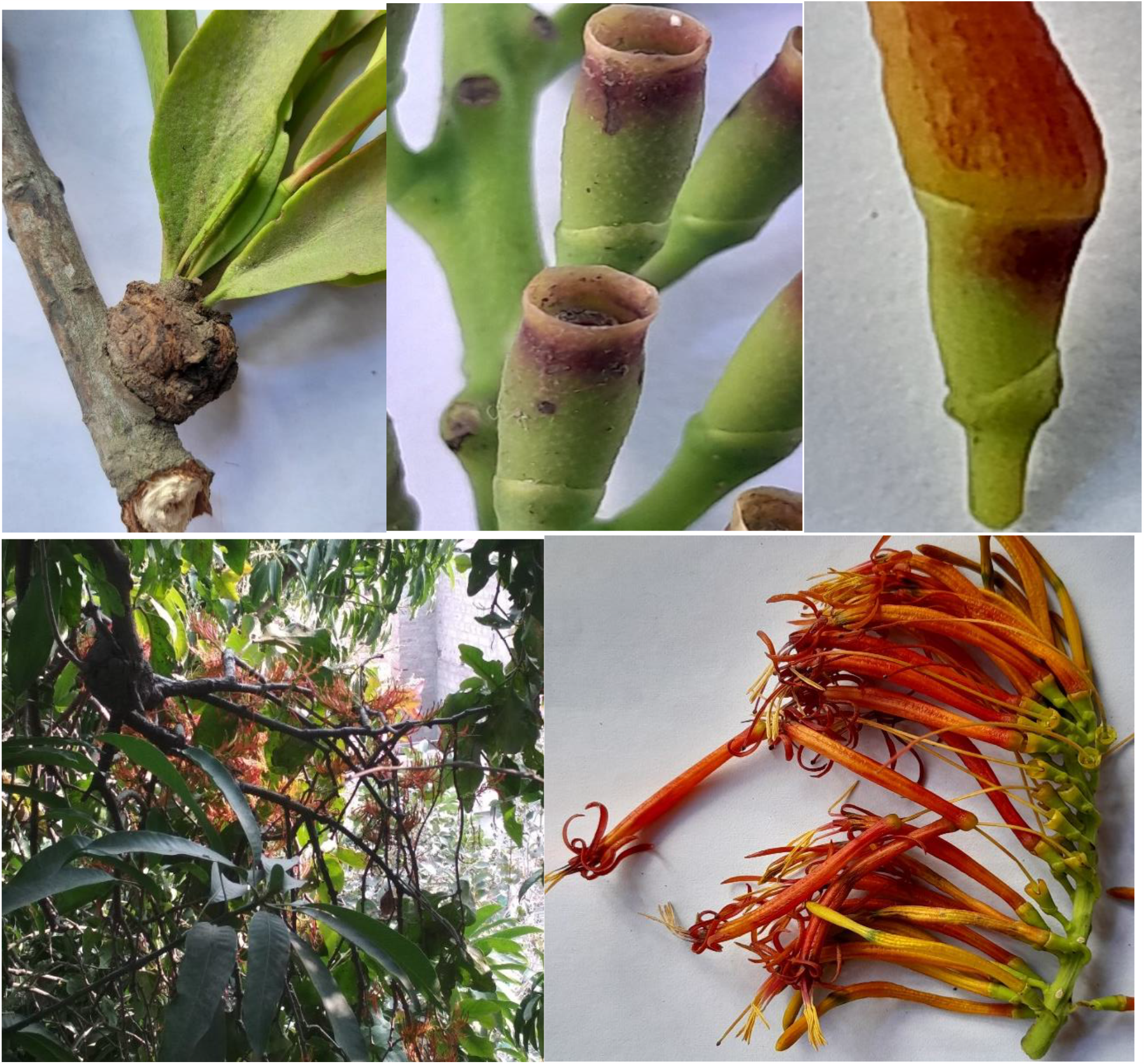
*Dendrophthoe falcata* (Clockwise from top left): A haustorium with branch; growing fruit; a calyculus with distinct bract; a bunch of mature flowers (corolla tube separated from calyculus); mistletoe with a distinct wood gall in the mango plant.

Stem shrubby, erect, round, solid, woody, smooth and dark grey in colour. Nodes are very distinct, in some cases swollen with which the petiole is attached. Branches dark greyish in colour. Epicortical runner attached with the host plant by means of haustoria which is also woody and corm-like from which leaves grow (Fig. 3 and 5).

**Fig. 4.**
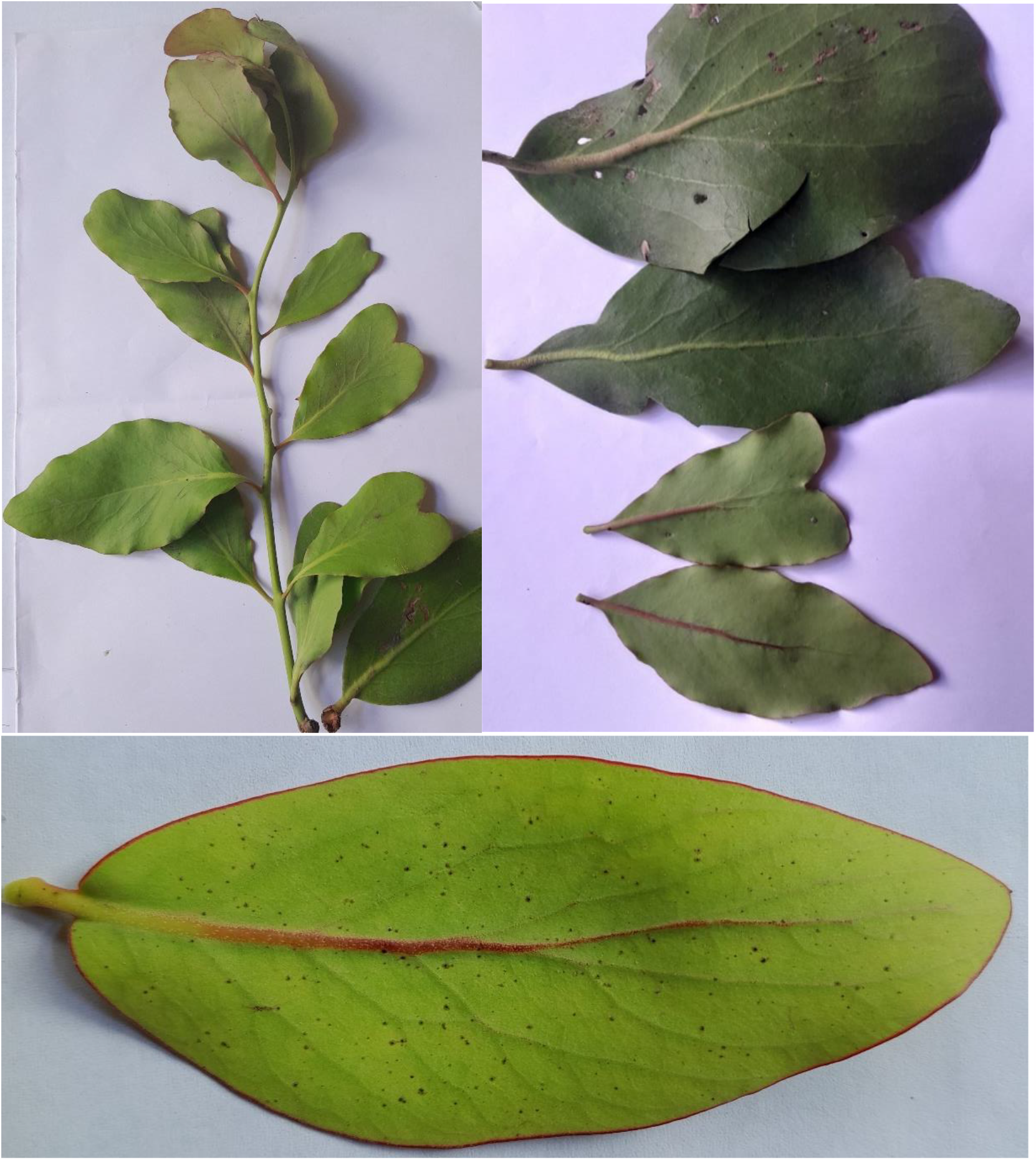
*Dendrophthoe falcata* (Clockwise from top left): A young branch showing phyllotaxy; different types of leaves; a young leaf with red margin and midrib (ventral side).

**Fig. 5.**
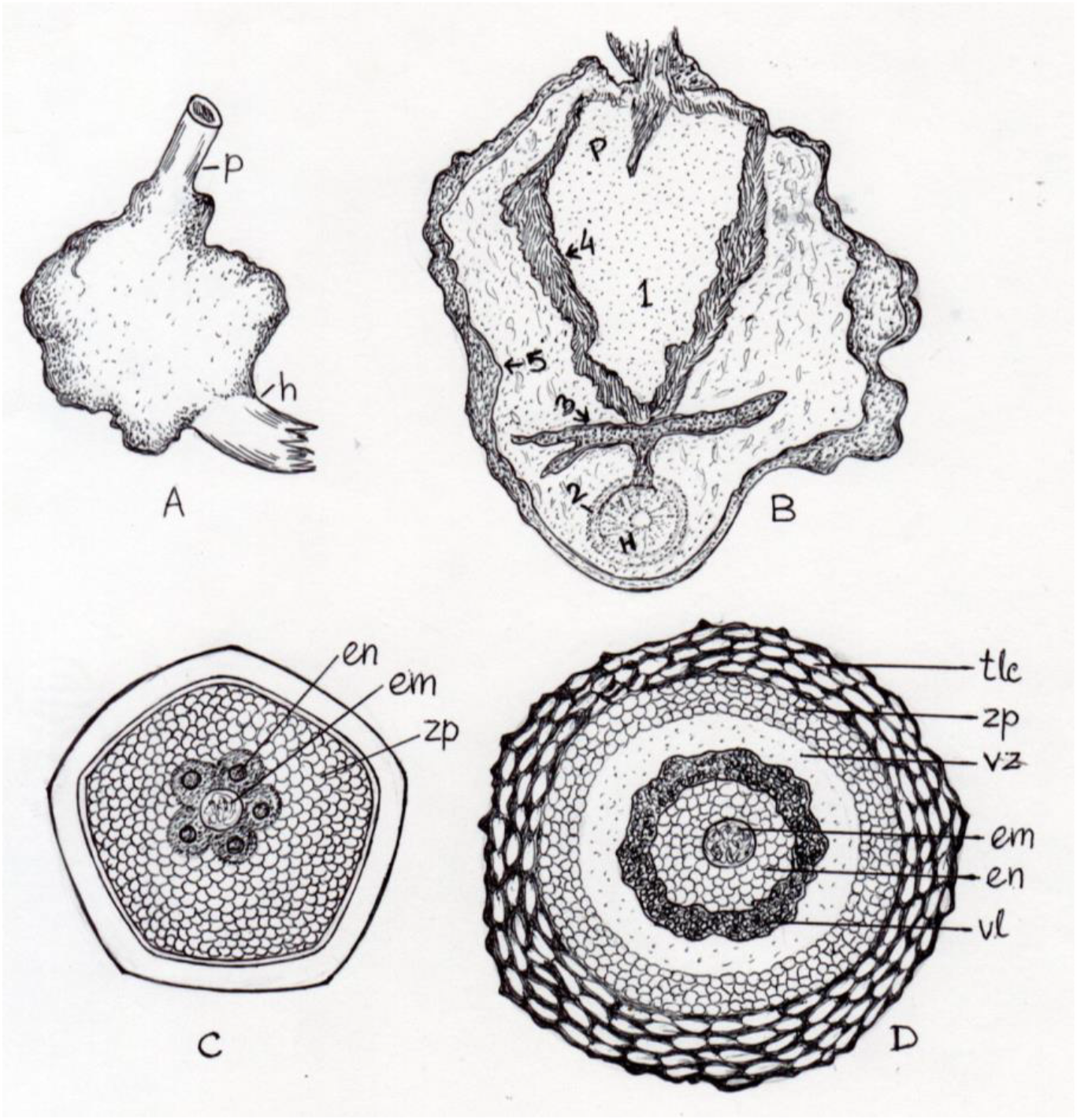
*Dendrophthoe falcata*. A. A haustorium, p-parasite, h-host; B. Section of haustorium, P-parasite, H-host, 1-penetration peg, 2-host cambium, 3-collapsed zone, 4-reddish brown secretion, 5-bark like dermal tissue of haustorium; C. Section of style, em-embryo, en-endosperm, zp-zone of parenchyma; D. Section of immature fruit, tlc-thick leathery coat, vz-vascular zone, vl-viscid layer.

Leaves opposite (mostly in young stem) or subopposite. Leaves are dark green, glabrous, thickly coriaceous. Petiole short, 5-8 mm. Lamina varies widely ovate, obovate, or elliptical; margin entire and undulate, cuneate or attenuate at the base. Apex also varies widely, may be acute (young leaf), obtuse or retuse (Fig. 4). In the young leaf, the leaf margin is with a distinct narrow red border and on the ventral surface, midrib is also pale red in colour (vs. green midrib on the dorsal surface) (Fig. 4). The size of the leaves also varies widely, 78-135 x 36-48 mm (in young leaf) to 140-160 x 70-94 mm (mature leaf). Venation pinnate with distinct midrib and the main laterals usually visible above and often more distinctly below.

Inflorescence raceme, axis 40-80 mm with 30-50 flowers. Pedicel short. Flowers are tubular with a green base. Flower is bisexual, actinomorphic, pentamerous, pedicellate, bracteate. Bract is small leaf-like, thick, and green, encircling half of the tubular calyculus and almost one-third in height of the calyculus (Fig. 6. I). Flower parts are arranged in acropetal succession: calyx (calyculus), corolla, stamens, and carpels (6. A).

**Fig. 6.**
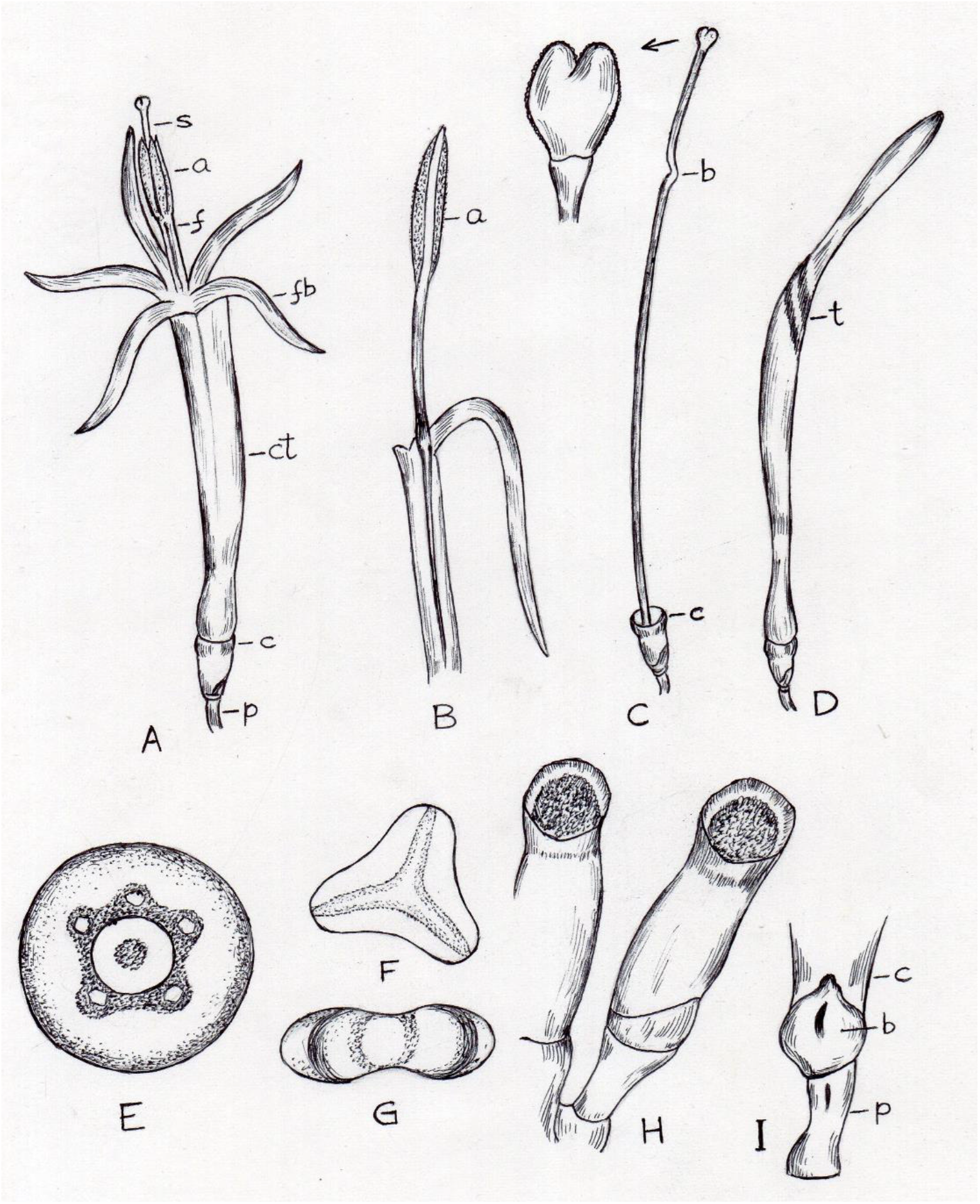
*Dendrophthoe falcata*. A. Complete flower, s-style, a-anther, f-filament, fb-reflex lobe, ct-corolla tube, c-calyculus/calyx, p-pedicel; B. Petaloid stamen, a-anther; C. Style, b-bend, c-calyculus/calyx; D. A mature flower bud, t-throat; E. Section of ovary; F. A pollen (polar view); G. A pollen (equatorial view); H. Immature fruit; I. Lower part of a flower, c-calyculus/calyx, b-bract, p-pedicel.

Calyx (Calyculus): Sepals fused to form a tubular epigynous rim often called “calyculus” whose morphological nature still in dispute. The distal one-third part of the tube is thin with entire margin, weakly pentagonal and pale red in colour (Fig. 2 and 6. C).

Corolla: In open flower, the petals are fused to form a corolla tube and five free anterior lobes are reflexed after anthesis. Corolla tube is long, dark red in colour and more than double in length compared to reflex lobes. The tube is narrow at the base and the basal globular part is pale red in colour. Reflex lobes are linear in shape and yellowish to red in colour on maturity (Fig. 2). Aestivation valvate.

The mature flower bud is long, uniformly widened upwards, slightly narrow at the base, weakly clavate and acute at the apex. The junction of the corolla tube and reflex lobes *i. e*. throat is swollen and green in colour. Usually, reflex lobular part is one-third in length of the whole bud. In immature flower bud, the upper part of the throat is greenish yellow in colour and bent (30°-40° of the axis of corolla tube) (Fig. 2 and 6. D).

Androecium: Stamens are five in number (pentandrous) and epipetalous (Fig. 6. B). Anthers form a temporary syngeny (not coalesces with each other) through which style extended (Fig. 6. A). Attachment of anther adnate (two anther lobes are connected longitudinally on both sides of the connective) and dehiscence of anthers longitudinal. Anthers acute at the apex. Anther lobes are equal or slightly longer than the free part of the filaments. Filaments are deep red in colour though anther lobes are yellow. Lower part of each filament is also red in colour, fuse with nearly the entire length of the corolla tube (Fig. 6. B).

Gynoecium: Monocarpellary pistil. Ovary is small and within calyx. Style is very long, filiform extending beyond the anther lobes. Style is apical in position and weakly pentagonal. Stigma bifid (Fig. 6. C). The colour of the style changes on maturity (yellowish green to dark red). Moreover, anterior cf. one-fifth part of the style is either dark in colour (in immature style) or with a bend in mature stage (Fig. 2, and 6. C). The ovary is inferior, 1 celled, without placenta, sporogenous cells in a single central mass in the ovarian cavity (Fig. 6. E). Section of style shows a central embryo encircled by growing endosperm (Fig. 5. C).

Pollen unit: Monad; Dispersal unit: Monad; Size: 0.22-0.25μm; Shape: Concave triangular (polar view), Emarginate (equatorial view); P/E ratio: Oblate; Sculpturing: Psitate; Aperture number: 3 (Fig. 6. F and G).

Fruit: Fruit 1-seeded berry with calyx remnant persistent at the apex, size 13-13.5 x 5.5-6 mm and prolate ellipsoid in shape. Seed solitary, spheroid, with five finger like processes at one side and opposite side with a small white, spongy, lobed cap - caruncle. The whole seed is within a white viscid, rubbery, mucilaginous mesocarp. Endosperm is white in colour with a protruding green globular radicle. Embryo green, about 5-6 mm long, cotyledons fused together, a globular radicle at the apex (Fig. xx). Section of young fruit shows a central embryo encircled by endosperm which is covered by a thick viscid layer, a salient feature of the family, Loranthaceae. The outermost layer is thick and leathery (Fig.5. D).

Floral measurements (in mm): Pedicel 2.7-3.1; bract 1.5-1.8; flower bud 38-40; calyx or calyculus 5.0-5.2; corolla tube 28.5-31.5, reflex lobe 13-14; free filament 5.4-6.8, anther 6.0-6.2; ovary 3.5-4.0, style 43-44.5, stigma 1-1.

Flowering: December to February.

Mistletoe status in *Mangifera indica*: Robust

In the section of root, a thick layer of periderm followed by a layer of phloem is present. Hexagonal xylem vessels are arranged in circular fashion, the number of which varies from 26 to 28. A layer of vascular cambium is present in between phloem and xylem (Fig. 7E).

**Fig. 7.**
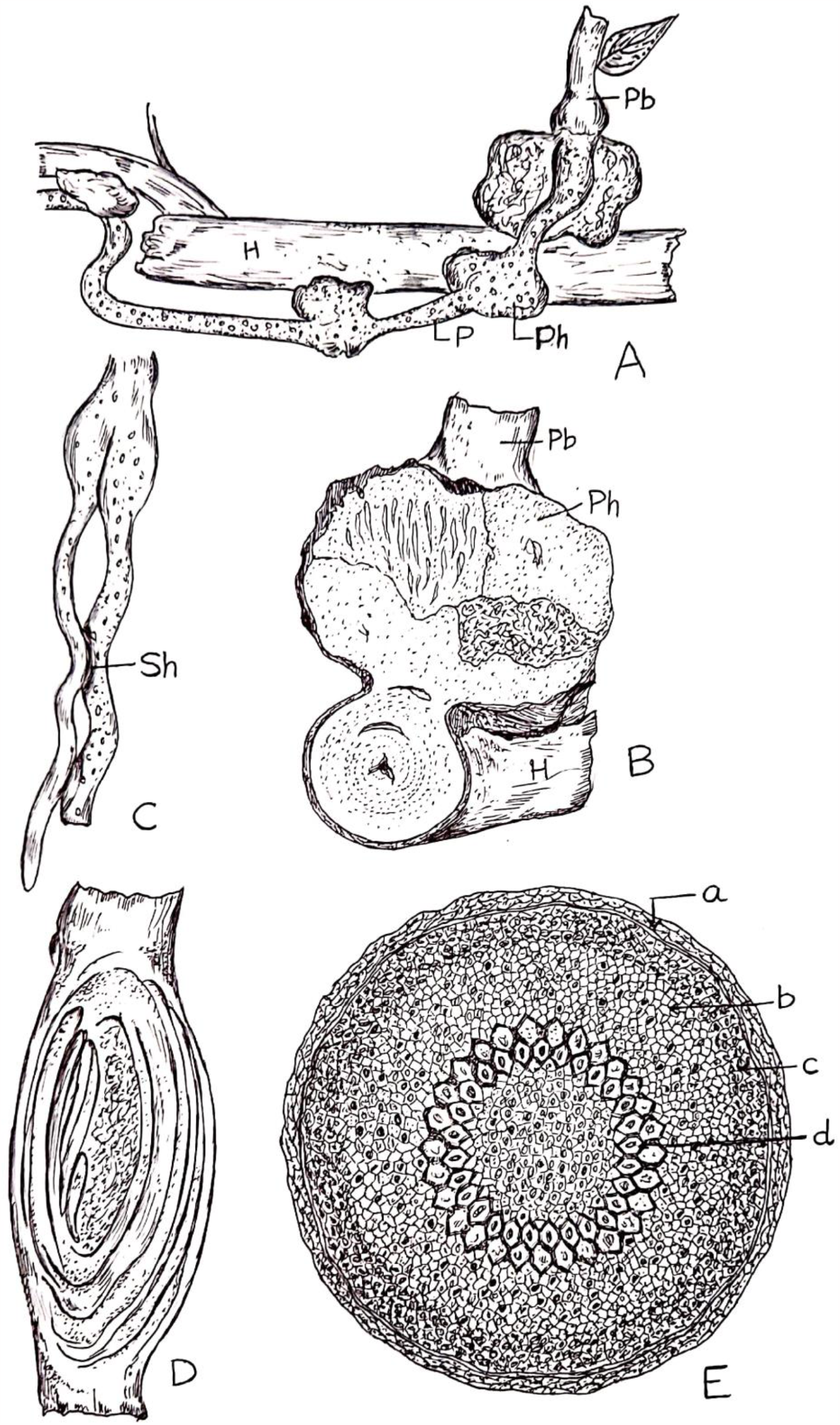
*Dendrophthoe falcata*: A. Multiple haustoria, H-host, P-parasite, Pb-parasitic branch, Ph-primary haustorium; B. Secondary haustorium (Sh) in between two growing roots; C. Section of haustorium; D. Structure of secondary haustorium; E. Section of root, a-periderm, b-vascular cambium, c-phloem, d-xylem.

Haustorium: I observed three types of haustoria namely woody gall, clasping union and epicortical roots in three different species. In mango and citrus plant “woody gall”, a fusiform swelling is formed in the host parasitic connection. But in guava plant, “clasping union” type of haustorium is present. In this type, an enlargement of the parasite root partially encircles the host branch and flat type. Moreover, multiple haustoria run in parallel to the host branch forming “epicortical root” type of haustoria (Fig. 7A). In both citrus and mango plant, from large corm like woody gall haustoria parasitic branches grow and hanging in the air with inflorescence. But in guava plant, parasitic branches only develop when the haustorium become large and round (Fig. 7A, C). Stunted growth of the branch observed in both mango and citrus plant while in guava plant, growth retardation in the branch containing epicortical root is not distinct. In guava plant, haustorium are of two types, primary or true haustorium which penetrate the host stem to collect the water while another type, secondary haustorium only help to anchor the two parts of the parasite root (Fig. 7B). This secondary haustorium is hollow, oval in shape, with 5-6 concentric ring-like thickening at the periphery and never penetrates the other stem to suck water (Fig. 7D).

I also observed another interesting thing. Though a large mango tree is heavily infected with the mistletoe but other host plants like citrus and guava plants in the vicinity are not at all affected while guava plant remain unaffected in the proximity of the heavily infected citrus plant.

Section of the wood gall type of haustorium reveals that when the root of parasite (haustorium) encountered the host tissue secondary xylem is formed between the endophyte and the host cambium. The woody root form a distinct conical penetration peg (containing xylem) the anterior of which touches the host cambium. The reddish-brown outline of the penetration peg is the secretion of the parasitic xylem. The penetration peg is halted when it reaches the host cambium. At this point, the tip of the peg flattened and promotes further growth against the surface of the host secondary xylem resulting flanges. In this flattened portion, a collapsed zone with numerous black grains develops. These are made up of crushed cell walls from both partners. Haustoria tissue further develops around the penetration peg and form bark like dermal tissue of dark grey colour (Fig. 1 and 5. A, B).

## Discussion

Whether calyculus is homologous with the calyx is debatable. Patil and Pai (1984) on the basis of vascular traces designated this whorl as a true calyx, an agreement with Agarwal (1963) and Smith and Smith (1942). Modern workers (Stanffer 1961a, Patil and Pai 1984, Endress 1994, Kuijit 2013, Der and Nickrent 2008) also regard this as a remnant of calyx. Though another school advocate the calyx as a bracteolar origin (Venkata Rao 1964, Wanntorp & Ronse De Craene 2009) and proposed the “Biseriate corolla concept” for Loranthus (Wanntorp & Ronse De Craene 2009) which is strongly criticized by Kuijt (2013). To avoid controversy, I consider the whorl as calyx.

In the present study, the colour difference (in young stage) or bend (in mature stage) in the style of mistletoe is a very peculiar feature. Singh (1952) reported a conspicuous bend and colour difference in the style of *Dedrophthoe falcata*. The upper part of the bend is with reddish tint while the lower part is yellow. He also noted that several embryo sacs develop simultaneously in the ovary and their tips, enclosing the egg apparatus, extend up to two-thirds the length of the style. In the present study, the colour difference is distinct in young style, deep green above and pale green or yellow below while in mature style, bend is distinct without any conspicuous colour difference. A bend in the style of *Dendrophthoe* can be explained with the findings of Johri and Bhatnagar (1972) and Johri (1984). In Loranthaceae, the extension of embryo sac to various heights in the style is a significant feature (Johri, 2012). Johri and Bhatnagar (1972) reported that in *Dendrophthoe*, the embryo sac reaches almost 75% of the style. Based on these morphological and embryological peculiarities of *D. falcata*, Singh (1952) commented that the lower part below the bend belongs to ovary while the distal upper part represents true style. The present observation also supports this hypothesis.

There is a controversy regarding the presence or absence of placenta in Loranthaceae. Griffith (1838, 1844) reported free central placenta in two species of *Loranthus*. Later, Schaeppi and Steindl (1942) studied nine genera and 121 species of Loranthoideae and reported that except a few, there is no placenta including *D. pentandra*. In *D. falcata* (present study), there is neither an ovule nor a placenta which endorses the observation of Singh (1952).

Pollen features of *D. falcata* in the present study is very similar to that of Heigh (2021). Like all Lorantheae pollen, pollen of *D. falcata* is of Type B (Grimsson et al. 2018).

Like *D. falcata*, seeds of *D. curvata* also bears five finger like processes (Barlow 1995). Probably these finger-like processes or fimbriae play a significant role in attachment of the seed to the host stem. At first, the seeds glued to the stem surface by the viscid mucilage, then fimbriae help in anchoring the seed with the surface tightly and finally the radicle grows and penetrate the bark after which mesocarp along with caruncle is shed off.

In woody hemiparasite plant like *D. falcata*, thickening and strengthening of the root system is necessary in supporting the trunk. For this reason, secondary vascular cambium is developed in the epicortical root and periderm forms due to extensive secondary growth which render mechanical support to the root.

Woody gall formation in mango tree is a fusiform swelling of the host-parasite connection. Local swelling of the host branch associated with woody gall formation is not only entirely caused by host branch but also due to proliferation of parasitic endophytic tissue within the host wood. Similar phenomena have observed in other woody gall forming mistletoes (Srivastava & Esau 1961, Hamilton & Barlow 1963, Calvin & Wilson 1998, Teixeira-Costa 2015).

Anatomy of the host-parasite interface in *D. falcata* is quite similar to all reviewed root-parasitic within Santalales (Fineran & Hocking 1983). Host-haustorium interaction at tissue level is observed by several authors (Kuijt 1965, Fineram 1965, Pate et al. 1990, Calvin and Wilson 1998, Teixeira-Costa 2021) and all of them submitted plausible explanations of how the root parasite absorb water and minerals from the host xylem. Kuijt (1965) reported absence of phloem in Santalalean haustoria and the vascular contact with host is exclusively xylary. In this order, haustorial xylem consists of reticulate-pitted vessel members in a matrix of parenchyma. Fineram (1965), Kuijt (1965), Pate et al. (1990) reported the growth of haustoria in host tissue in a very similar pattern as observed in the present study.

Toth and Kujit (1977) studied the anatomy and ultrastructure of haustorium in a species of Santalaceae. They observed dark staining non-cellular layer at the junction of endophyte and host tissue, that seems to be made up of crushed cell walls from both partners and there is no cytoplasmic connection between host and parasite. Fineram (1985) in his review, mentioned that graniferous tracheary elements, an unusual xylem conducting cells, is characterized by having structural material in the lumen. These elements are present typically in the body of haustorium of root parasite especially in the expanded xylem tissue. These graniferous elements are present in all species of Santalaceae, two species of root parasite Loranthaceae and Lauraceae (Rajanna and Shivamurthy 2001). These grains are of different nature, either starch or protein. It was suggested that in Santalaceae, it might be a device for regulating the flow of xylem sap through the haustorium. Fineram and Bullock (1979) and Joel (2013) also opined that this xylem conduit help to regulate sap flow from host to parasite. Heide-Jorgensen (2008) commented that these granules help in lowering haustorium osmotic potential and thus prevent reverse sap flux from the parasite to the host. In the present study, at the junction of extended endophyte and host tissue, there is a lumen containing numerous black grains and no direct tissue connection in between the two partners. Through this structure water conduction may takes place from host to parasite in apoplastic pathway.

Danser (1931, 1938) rightly pointed out that *D. falcata* is very difficult to circumscribe. The species integrades with other species in different geographical localities. As for example, the floral and leaf characters are very similar to *D. pentrada* and *D. praelonga* in Malesia and to *D. vitellina* in northern Australia. Moreover, in New Guinea, *D. gjellerupii* can be distinguished from *D. falcata* only on corolla dimensions. The morphological variation of characters in different geographical localities indicate that *D. falcata* is a polymorphic species having wide range of distribution. Recent discoveries of new species of *Dendrophthoe* especially from India is doubtful (Singh et al. 2020b, Sivaramkrishna 2021). Because the species are described on the basis of a few overlapping morphological diagnostic characters and comparisons are made only with a nearby species.

## Acknowledgement

I am deeply indebted to my students Sahab and Avijit and my neighbour Mohan for collection of the mistletoe plant.

## Notes

### Competing Interest Statement

The authors have declared no competing interest.

